# DNA Chisel, a versatile sequence optimizer

**DOI:** 10.1101/2019.12.16.877480

**Authors:** Valentin Zulkower, Susan Rosser

## Abstract

**Motivation:** Accounting for biological and practical requirements in DNA sequence design often results in challenging optimization problems. Current software solutions are problem-specific and hard to combine.

**Results:** DNA Chisel is an easy-to-use, easy-to-extend sequence optimization framework allowing to freely define and combine optimization specifications via Python scripts or Genbank annotations.

**Availability:** as a web application (https://cuba.genomefoundry.org/sculpt_a_sequence) or open-source Python library (code and documentation at https://github.com/Edinburgh-Genome-Foundry/DNAChisel).

**Supplementary information:** attached.

## 1 Introduction

Advances in DNA synthesis have made it possible for biologists to routinely order DNA constructs with custom nucleotide sequences (Kosuri and Church, 2014). While a sequence’s primary purpose may be the study or engineering of an organism, its design may also account for manufacturability, host compatibility, and other practical requirements, resulting in complex multi-constrained optimization problems.

Software solutions have been proposed to address various scenarios, including host-specific codon optimization or harmonization (Richardson *et al.*, 2012; Claassens *et al.*, 2017), gene expression enhancement via CpG island enrichment (Raab *et al.*, 2010), the design of biologically neutral sequences (Casini *et al.*, 2014), or the removal of synthesis-impeding DNA patterns (Oberortner *et al.*, 2017). However, these projects focus on specific objectives and predetermined sequence locations (such as coding regions), and are hard to integrate in a same workflow, as their optimizations may undo one another. The D-tailor framework (Guimaraes *et al.*, 2014) introduced the possibility to combine different specifications via Python scripts, with a focus on the exploration of multi-objective problems.

DNA Chisel improves on these approaches with new methods for constraints resolution and objective maximization, the possibility for researchers to provide design specifications via Genbank annotations, support for circular sequences, detailed output reports, and over 15 built-in classes of specifications (listed in Supplementary Section 1A) which can be freely composed to handle any combination of the optimization objectives mentioned above, and extended with user-defined specifications. The framework can be used as a web application, a Python library, or a command-line application, making it suitable for a wide range of uses.

## 2 Problem definition

An optimization problem is defined in DNA Chisel by a list of global or local specifications against which a starting sequence will be optimized. A specification can be either a hard constraint, which must be satisfied in the final sequence, or an optimization objective, whose score must be maximized. For instance, specification `AvoidChanges` can be used as a constraint to forbid sequence modifications in a given region, or as an objective to simply penalize changes in that region. In presence of multiple optimization objectives, which can be attributed relative weights, the total weighted score is maximized.

A problem can be defined by annotating a Genbank record to indicate the nature and scope of the different specifications to be applied to the record’s sequence, as shown in Figure 1A. This can be done interactively using a free Genbank editing software such as SnapGene Viewer (www.snapgene.com) or Benchling (www.benchling.com). Annotated Genbank files can be uploaded to a public web application (Figure 1B), which will optimize the sequence and return a multi-file report (see Supplementary Information 2 for an example). The report features the optimized sequence in Genbank format with annotations indicating modified nucleotides, a PDF report summarizing the changes (as shown in Figure 1C), and in case of unsuccessful constraint resolution, a troubleshooting figure representing the problematic region.

**Fig. 1.**
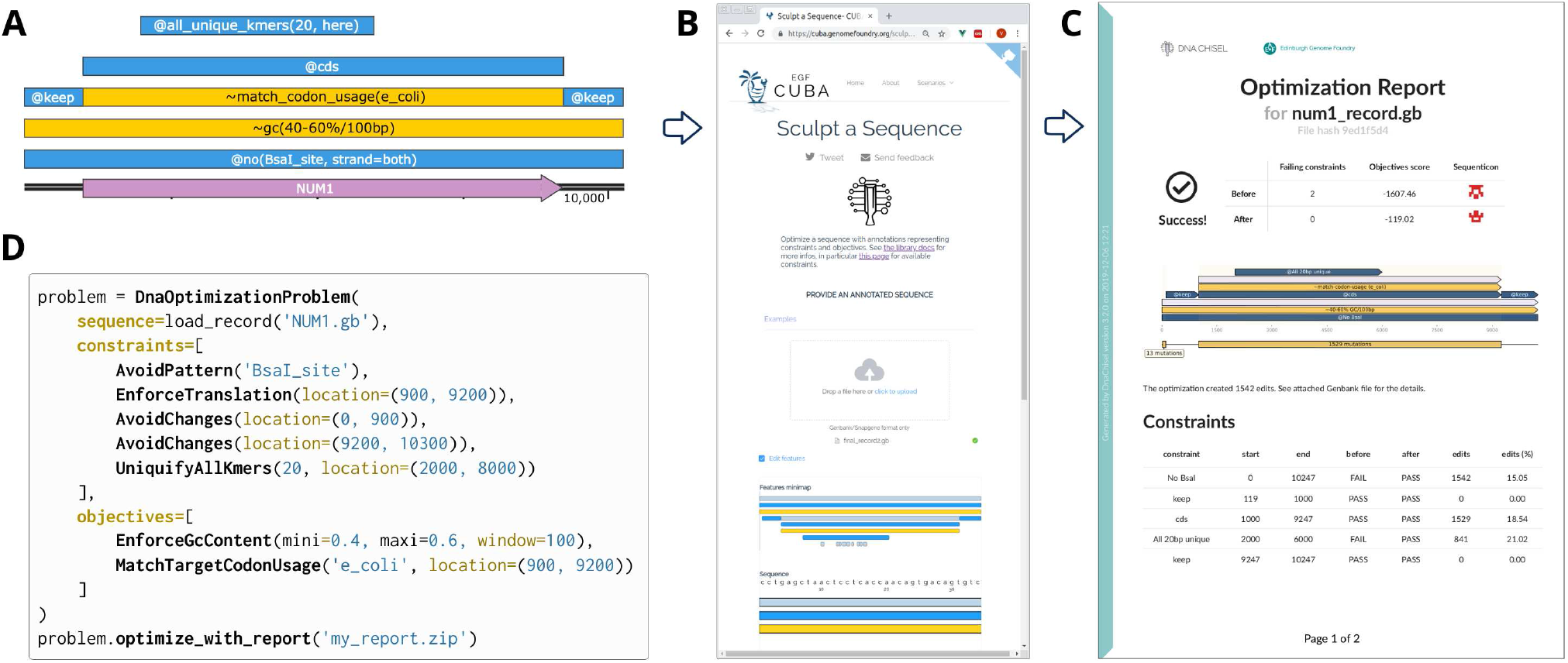
Using DNA Chisel online and via scripts. (A) Sequence optimization problem defined via Genbank annotations. Prefix “∼” in annotations indicate optimization objectives, while “@” denotes hard constraints. Gene NUM1 of S. cerevisiae will be codon optimized for E. coli, and have a GC content of 40-60% on every 100-nucleotide window. Its flanking promoter and terminator regions are protected against changes by @keep annotations(shorthand for @AvoidChanges). BsaI restriction sites will be removed (@no(BsaI_site) constraint) and undesirable homologies occurring naturally in the gene sequence will be removed in the region marked @all_unique_kmers, using codon-synonymous mutations only as enforced by the @cds constraint. (B) Screenshot of the online DNA Chisel application featuring a file dropzone and a basic Genbank annotations editor. (C) PDF report returned by DNA Chisel to summarize the problem’s specifications and highlight constraints resolutions and objectives score improvements. (D) Python script defining the same optimization problem as in panel A.

Optimization problems can also be defined and resolved via Python scripts (Figure 1D), enabling the programmatic definition of complex optimization scenarios involving hundreds of specifications. Supplementary Sections 1B and 1C provide example scripts for mass sequence pattern removal, and genome-scale gene domestication.

## 3 Implementation

DNA Chisel is implemented in Python. It relies on the Biopython library (Cock *et al.*, 2009) for Genbank operations and restriction enzyme data, and on the Codon Usage Database (Nakamura, 2000) for codon usage data.

The optimization algorithm, described in more detail in Supplementary Section 1D, has two main steps: (1) resolution of all hard constraints, ignoring optimization objectives, and (2) objectives maximization with respect to the constraints. This steps separation allows to detect incompatible constraints early, and quickly reach satisfactory solutions.

During each step, unsatisfactory sequences regions are detected and separately optimized. The solver speeds up region optimization by creating a local version of the problem with simplified specifications locally equivalent to the original problem’s specifications. A region’s sequence is optimized via either random mutations or an exhaustive search through all possible sequence variants, depending on the number of variants. Specifications can also implement custom local resolution methods to improve the solver’s efficiency. For instance, an `insert(CGTCTC)` constraint will attempt to place the pattern “CGTCTC” at different locations of the sequence, rather than relying on random or exhaustive search to create the pattern in the sequence.

## 4 Framework extension with new specifications

The emergence of new biological applications often comes with new DNA sequence optimization requirements. In this perspective, DNA Chisel allows to define any new sequence specification that the Python language can express, without modifying the library’s code nor compromising the solver’s efficiency. A new specification is defined by creating a new Python class with custom sequence evaluation and local resolution methods, and the class can optionally be registered with DNA Chisel’s Genbank record parser to enable its use via Genbank annotations.

## Acknowledgements and Funding

We thank Ye Chang, Paulina Kanigowska, Simone Pignotti and Li Xing for useful discussions on this project. The Edinburgh Genome Foundry is supported by the BBSRC (BB/M025659/1, BB/M025640/1, and BB/M00029X/1 to SR) and the BBSRC/MRC/EPSRC funded UK Centre for Mammalian Synthetic Biology (BB/M0101804/1 to SR) as part of the RCUK’s Synthetic Biology for Growth programme.

## Supplementary Information

### A. Built-in specifications

The table below lists some specifications available in DNA Chisel v3.1 (please refer to the online documentation at https://edinburgh-genome-foundry.github.io/DnaChisel for an up-to-date list). For each specification class, we provide the following information:

- **Specification class:** name of the specification for use in Python scripts.
- **Example of annotation:** example label to use when annotating genbank records. Note that shorthand annotations can be used, e.g. @gc(39%) instead of `EnforceGCContent(target=0.39)` (both annotations will be recognized by DNA Chisel)
- **Effect:** short description of the specification’s effect on a sequence.
- **Examples of use:** scenarios in which the specification can be relevant.

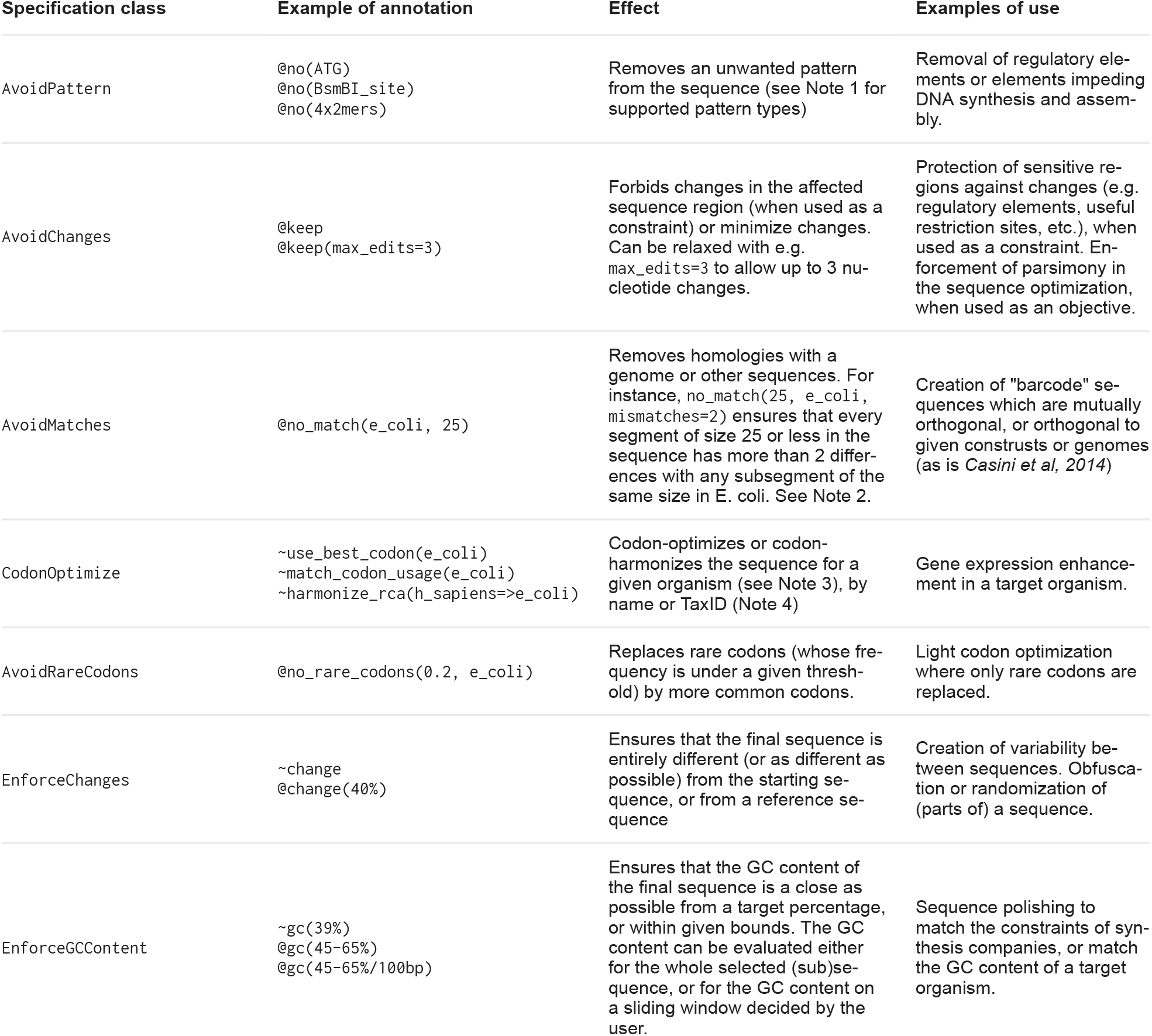
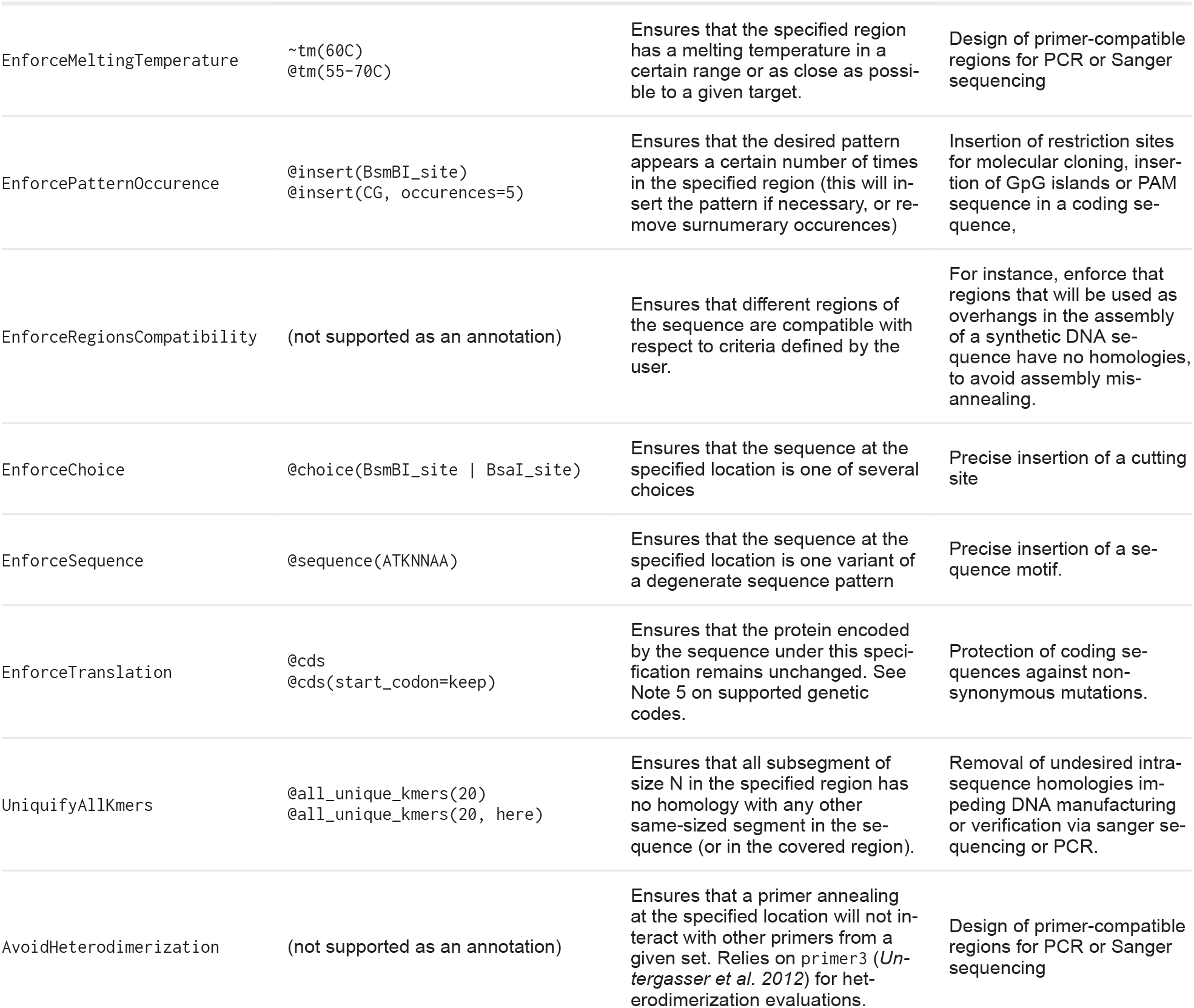

#### Notes

1. Patterns provided to specifications such as `AvoidPattern` and `EnforcePattern` can be either:

- A DNA sequence (e.g. `ATGG`)
- A degenerate DNA sequence (e.g. `ATH` to represent `ATC` *or* `ATT`)
- An enzyme restriction site (using the annotation `BsmBI_site`) or any other built-in DNA Chisel pattern class (e.g. `9×A` to represent a homopolymer with 9 “A”s in a row)
- A Position-Specific Scoring Matrix (PSSM), representing a DNA motif (*Stormo et al. 1982*). Such patterns are not supported for use via genbank annotations and can only be defined when using DNA Chisel via Python scripts.
- A regular expression.
2. Specification `AvoidMatches` relies on the Bowtie software (*Langmead et al. 2009*) to find perfect (or near-perfect) matches between short subsegments of a sequence and a Bowtie index. As of v3.0 of DNA Chisel, another specification, `AvoidBlastMatches`, allows to remove any match found using the BLAST algorithm instead of Bowtie, allowing for more relaxed dissimilarity in the alignments. Note that `AvoidMatches` normally takes a path to a Bowtie index as an argument, and can only be used as shown in the example (with `e_coli` as input) if the name `e_coli` has been associtated with a Bowtie index in the server.
3. Three optimization methods, with different objectives, are supported by the `CodonOptimize` specification in DNA Chisel v3.1:

- Replace each codon by the most frequent codon in the target organism. More precisely, the Codon Adaptiveness Index is maximized, as defined in *Sharp et al. 1987*.
- Ensure that the relative frequencies of the various codons of the sequence match the global codon usage of the target organism. Note that this objective, proposed as early as in *Hale and Thompson 1998*, appears throughout literature under different names, such as Individual Codon Usage Optimization (*Chung et al. 2012*), Global CAI Harmonization (*Mignon et al. 2018*), and Codon Harmonization (*Jayaral 2005*) - not to be confused with the more accepted codon harmonization procedure described next.
- Codon-harmonize the sequence by replacing each rare codon for the native organism by a rare synonymous codons in the target organism, and common codons by common codons. The score maximized relying on Relative Codon Adaptiveness, as proposed in *Claassens et. al, 2017* as a variant of the codon harmonization method first described in *Angov et al. 2008*.
4. DNA Chisel supports codon optimization for virtually any possible target organism thanks to the `python_codon_tables` library (pypi.org/project/python_codon_tables), which was developed specifically to support DNA Chisel and similar projects. Organisms can be specified using either a pre-registered organism name of `python_codon_tables` such as `e_coli`, `h_sapiens`, etc., or a taxonomic identifier, e.g. “6239”, corresponding to a codon usage table in the Codon Usage Database (kazusa.or.jp, *Nakamura et al. 2000*), at which case the codons usage table will be automatically downloaded from the database.
5. Any genetic code supported in Biopython’s CodonTable module is also supported in DNA Chisel. This includes the standard genetic code, the bacterial genetic code, the mithocondrial genetic code, and a dozen others. Note that for genetic code accepting a non-ATG start codon (e.g. GTG in bacteria) the user must provide a policy for the replacement the original coding sequence’s start codons, to either keep the codobn’s original sequence, always replace its sequence with ATG, or leave a choice of valid start-codon sequences.

### B. Example: generation of a restriction-free sequence

Several molecular biology applications require to remove restriction sites from a DNA sequence, for instance to allow the future assembly of the sequence with other DNA fragments using restriction-based cloning methods.

To demonstrate DNA Chisel’s ability to cope with a high number of such constraints, the following Python first generates a list of constraints forbidding the restrictions sites of 458 different enzymes, corresponding to all 6-cutter listed in the Biopython library.

The script then generates a 5000bp random sequence, which initially features ~1500 chance occurences of the restriction sites at various locations, and proceeds to optimize the sequence to remove all sites. It completes in 5 to 6 seconds on a Intel® Core™ i5-6500 CPU @ 3.20GHz × 4)

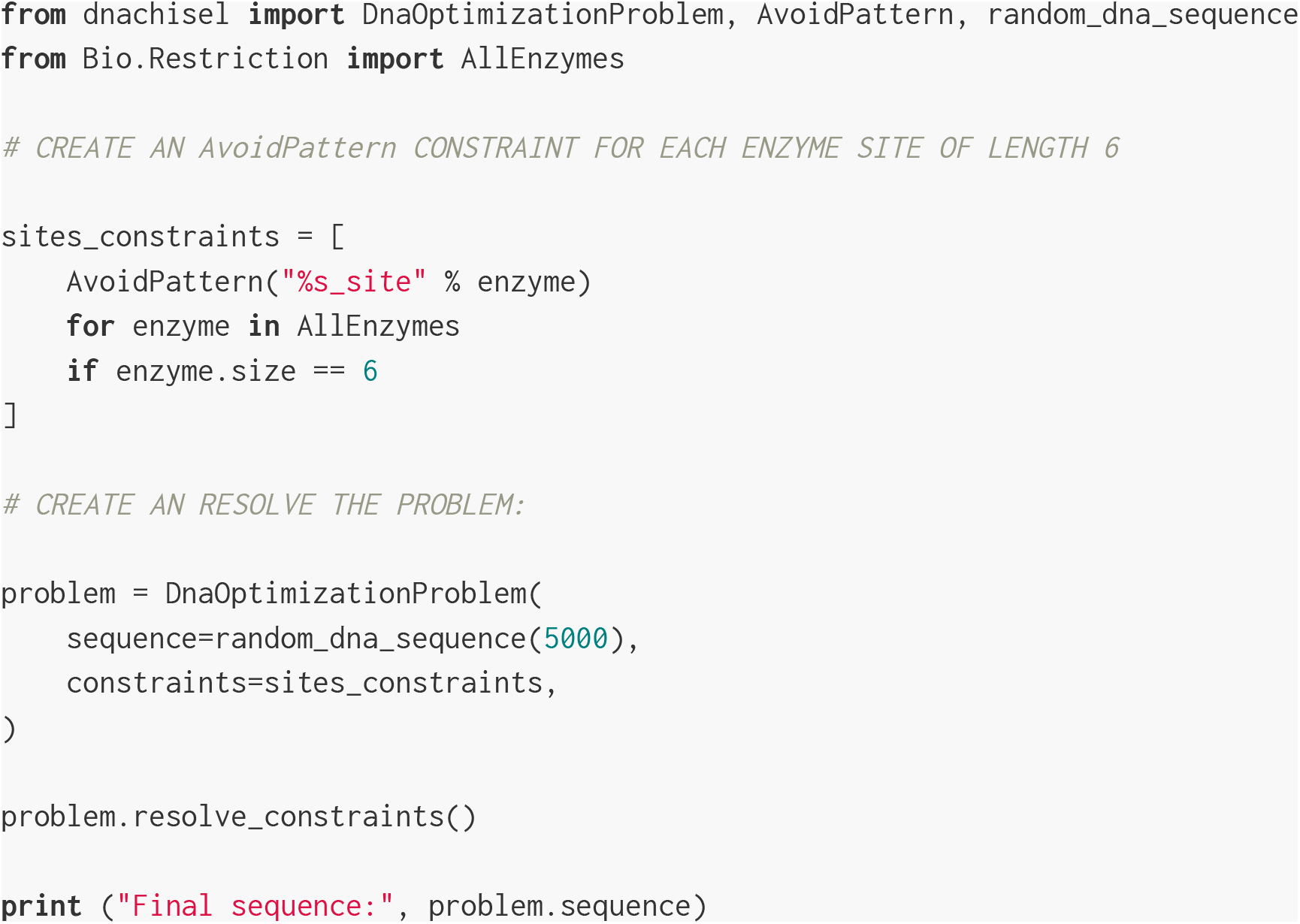

### C. Example: genome-wide gene optimization

The script below downloads the ~4Mbp *E. coli* genome and extracts its ~5000 genes one by one to optimize them (removal of BsaI and BsmBI enzyme sites impeding DNA assembly via Golden Gate assembly, and codon optimization for CAI index maximization) and write them in separate files (a scenario inspired by *Scher et al. 2019*). The script takes 5 to 6 minutes to run on an Intel® Core™ i5-6500 CPU, 3.20GHz × 4 (i.e. ~15 genes are optimized per second).

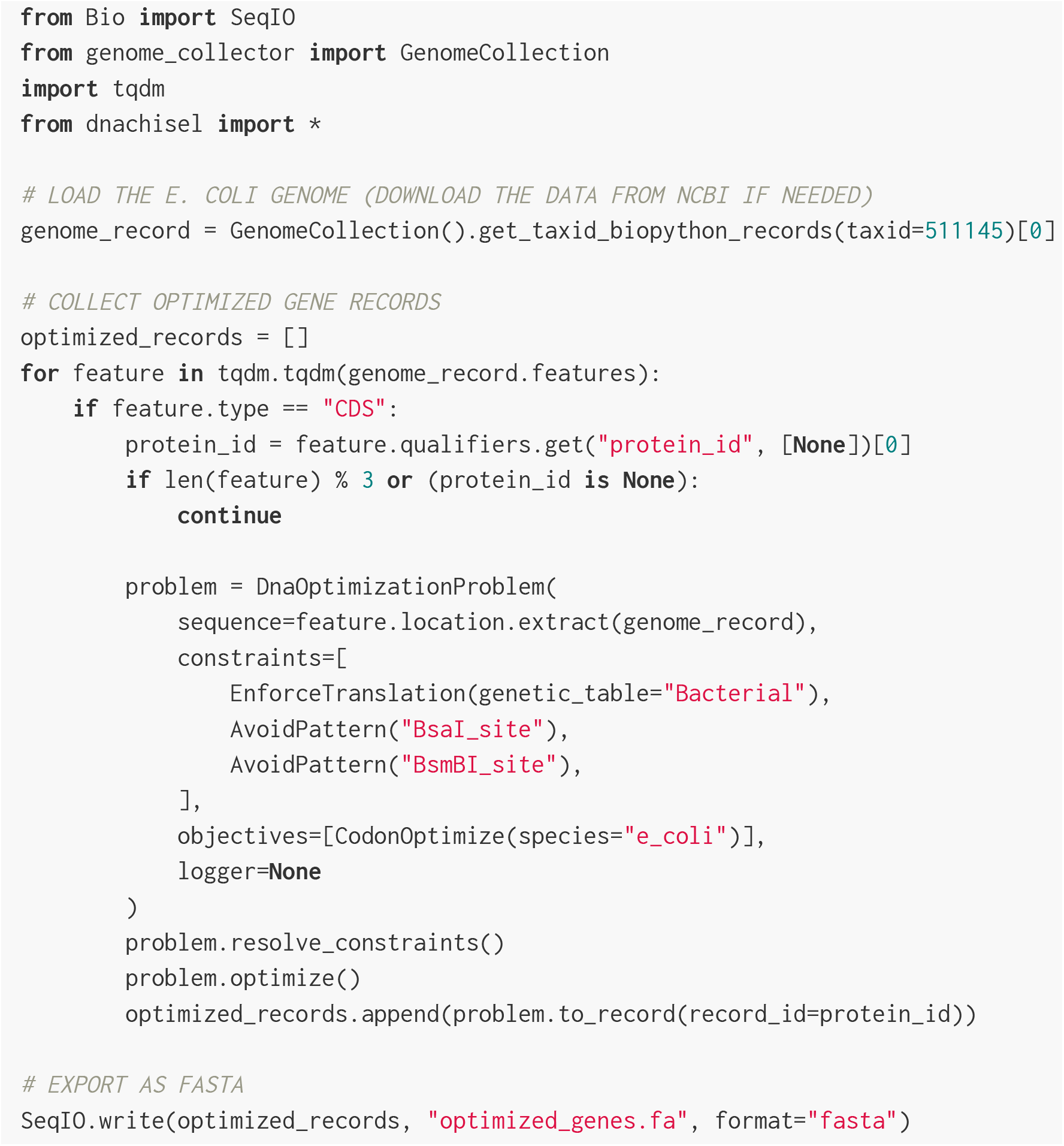

### D. Sequence optimization techniques in DNA Chisel

This section describes the mechanisms underlying DNA Chisel’s constraint resolution and objective optimization algorithms.

#### Mutation space

DNA Chisel relies on a “mutation space” data structure to guide sequence modifications during sequence optimization. The mutation space indicates the acceptable values for the different nucleotides and/or sub-segments of the sequence, and is determined by the problem’s nucleotide-restricting constraints. In an unconstrained problem, each nucleotide can be mutated to either A, T, G, or C. When an `EnforceTranslation` constraint is applied to a stop codon, it constrains the nucleotides triplet to only 3 possible choices, TAG, TGA, and TAA (see Figure S1 for an illustration).

Computed only once at problem initialization, the mutation space reflects the restrictions of all nucleotide-restricting constraints at once, incidentally allowing the early detection of incompatible constraints (i.e. constraints with no compatible solutions at certain sequences locations). More advantageously, constraints which are entirely guaranteed by restrictions of the mutation space (such as `AvoidChanges` and `EnforceTranslation`) can be removed from the problem after mutation space initialization, thereby significantly accelerating the optimization procedure.

Note that preexisting frameworks also restrict mutations in their own way. The D-tailor framework introduces manipulations of the mutation space, which differ significantly from DNA Chisel’s as they happen within the optimization loop and constrain the location of possible mutations, rather than the choice of nucleotides. Frameworks BOOST, D-tailor, and GeneOptimizer all implement a form of mutation space restriction to ensure synonymous mutations in coding sequences, but only for this particular use case, while DNA Chisel will combine mutation space restrictions coming from constraints such as @keep, @cds, @change, @sequence(…), @no_rare_codons(…), etc.

**Figure S1:**
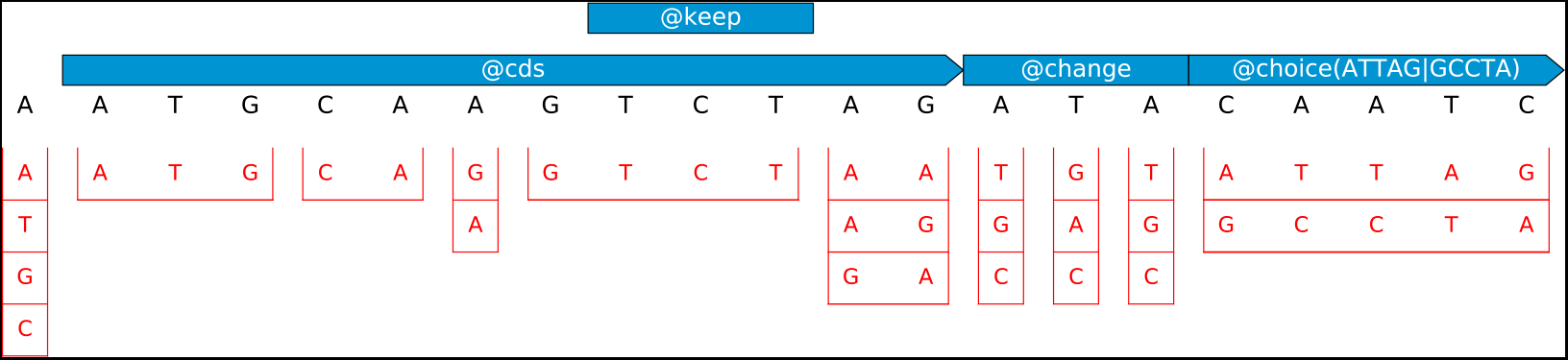
Restriction of the mutation space by different specification classes. In this example the problem consists of a 21-nucleotide sequence and four constraints. The mutation space, represented in red, consists of contiguous sub-segments, each associated with a set of sequence choices. For instance, the first nucleotide is unconstrained and can take any of the four possible values. The next nucleotides are constrained by @cds which enforces synonymous codons mutations, and these restrictions are combined with a @keep constraint which keeps the affected sequence segment in its original state.

#### Specification evaluation on a sequence

In DNA Chisel, each specification implements a custom evaluation method indicating how well a given sequence complies with the specification. The output is the same whether the specification is used as a hard constraint or an optimization objective, and comprises:

- A score indicating whether and how well the sequence complies with the specification (the more compliant, the higher the value). By convention, a negative score indicates that the specification is breached, while a score of 0 or more indicates compliance.
- A list of segments coordinates (start, end) indicating the locations of breaches (or, for optimization objectives, sub-optimal regions). For each breach, the coordinates indicate the largest sequence sub-segment where modifications are susceptible to resolve the breach.
- A message in plain English which can be used in reports to help users understand why the sequence does not comply with the specification.

#### Optimization algorithm overview

DNA Chisel’s algorithm first ensures that all constraints are verified, then optimizes the objectives with respect to the constraints. The solver follows the following procedure:

- **For each** constraint:

- Evaluate the constraint on the problem to find the location of all breaches.
- **For each** breach location, from left to right:

- Define a *local problem* to resolve the breach locally, while making sure not to create new breaches locally-verified constraints.
- **For each** optimization objective:

- Evaluate the objective on the problem to find the location of all sub-optimal regions.
- **For each** sub-optimal region, from left to right:

- Define a *local problem* and use it to optimize the local region in order to increase the overall objectives score while making sure that all constraints remain verified.

While some other frameworks also use local optimization, DNA Chisel introduces new techniques, described in the next paragraphs, to simplify local problems and accelerate their resolution.

#### Definition of local problems

In DNA Chisel, a local problem is a version of a problem where only a small segment of the sequence (denoted *[start, end]* in this section) will be mutated, in order to locally resolve a particular constraint breach, or increase the sequence’s fitness with respect to local objectives.

The mutation space of the local problem is obtained by *freezing* the problem’s mutation space outside of segment *[start, end]*, and the local problem’s specifications are a simplified version of the problem’s specifications using the following rules:

- Specifications whose location has no overlap with segment *[start, end]* won’t be affected by the local mutations, and can simply be removed from the local problem to avoid unnecessary evaluations.
- Any constraint that is not initially verified in the mutated region, except the constraint being resolved, is removed from the local problem (this constraint will be resolved in its own time in a subsequent iteration of the main “for each constraint” loop).
- Each remaining specification *S* can be simplified to a local version *S*′ whose evaluation is restricted to the neighborhood of *[start, end]* in order to detect any breach to *S* that can be caused by mutations on segment *[start, end]*. Said otherwise, the local specification *S*′ has a simpler and faster evaluation method, while being locally equivalent to *S*.

The computation of local forms for the specification (illustrated by an example in Figure S2) is one of the most computationally advantageous mechanisms introduced by DNA Chisel. Each specification class implements a custom localization method to ensure that the local specification will be as fast to compute as possible, while still being able to detect newly-create constraints breaches.

**Figure S2:**
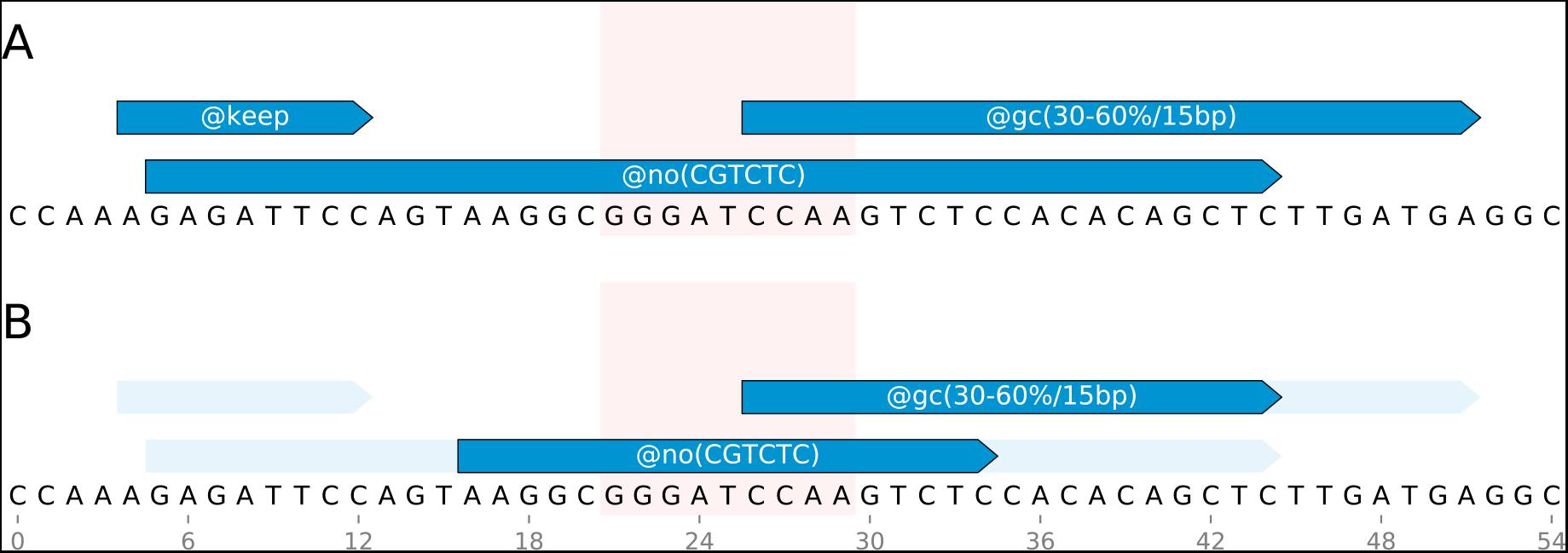
Problem simplification via localization. The problem represented in panel A is localized to the short segment *[21, 29]* (highlighted in red) to obtain the local problem represented in panel B. Note that the @keep specification disappears completely in the local problem as it does not overlap with the localization segment. Constraint @no(CGTCTC) is reduced to the a region corresponding to the localization segment with a 5-nucleotide addition on each flank, in order to capture any formation of new `CGTCTC` sites (notice that changing `A` to `C` at position 29 would create such a site at position 29-34). Constraint @gc(30-60%/15bp), which uses a 15bp window, gets reduced to the only 15bp windows that can be affected by changes in segment 21-29, i.e. reduced to segment 21-44.

#### Local sequence optimization

Given a local problem with mutations restricted to a sequence segment, DNA Chisel proceeds as follows:

1. Select a search method to explore the mutation space.
2. Use this method and find a sequence variant satisfying all constraints (when resolving constraints) or maximizing objectives scores (when optimizing objectives).
3. Replace the problem’s sequence with the successful variant, and move on to the next location to optimize.

In step 1, the search method is selected based on number of possible sequence variants in the local problem. This number can be easily computed from the mutation space, as the product of the number of choices at the different locations. For instance, in the problem shown in Figure S1, the mutation space allows

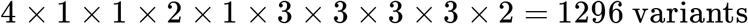

If the sequence admits only a small number of variants, the solver performs an *exhaustive search*. If this number is above a certain threshold (which can be set by the user, and defaults to 10,000), then a *guided random search* is used instead. Finally, in the (uncommon) case where the specification being resolved (or optimized) implements its own resolution method, this custom method will be used.

##### Exhaustive search

In an exhaustive search, the solver considers each possible sequence variants one after the other.

When the solver’s goal is the resolution of a constraint breach, the loop stops as soon as one variant verifying all constraints is found. If the loop completes without such variant being found, it can be concluded that the problem has no solution, as some constraints cannot be (simultaneously) satisfied. The solver then terminates and raises an error (when optimizing with a PDF report output, this error will be processed to generate a schema of the problematic region for troubleshooting).

When the solver aims at objectives score maximization, each variant is first evaluated against the problem’s constraints, and only constraints-complying variants are considered. The loop stops when a variant is found for which the total objective score equals the theoretical maximum (in the case where all objectives have an attributed theoretical maximum), or when all sequence variants have been evaluated, at which case the sequence variant with the highest score is retained.

Exhaustive searches present the advantage of being able to find even *rare* solutions (*needles in haystacks*), or providing evidence of the absence of solution. However, they can be slow for problems with a large mutation space, for which random searches are better adapted.

##### Guided random search

In a guided random search, the solver uses random mutations to iteratively progress towards a solution.

A sequence mutation is created by substituting the sequence of one sub-segment of the mutation space by another acceptable choice for this sub-segment. By default, 2 mutations are performed at each iteration, although this number can be set by the user. If the resulting *candidate sequence* performs better than the non-mutated sequence, it becomes the reference sequence for the next iterations, otherwise it is simply discarded, and the next iteration proposes new mutations of the current reference sequence. The maximum number of allowed iterations to reach a solution can also be set by the user, and defaults to 1000.

When the solver’s goal is the resolution of a constraint breach, each candidate sequence is evaluated against the local constraints. If all constraints are verified, the resolution stops and the candidate sequence is retained. Otherwise, the *constraints score* of the candidate sequence is computed by summing the scores of all failing constraints evaluations. If this score is above the score of the non-mutated sequence (meaning that the mutations made the sequence closer to being compliant), the candidate sequence becomes the reference sequence. If the maximum number of iterations is reached without finding a solution first, the solver terminates and returns an error. When optimizing with a PDF report output, the report will feature a schema of the problematic region. However, in this case, it is not to determine whether the failure was due to un-satisfiable constraints, or whether the solver simply failed to reach a solution in the allowed number of iterations.

When the solver aims at objectives score maximization, each candidate sequence is first evaluated against the local constraints, and if any breach is detected the candidate is directly discarded. Else, the sum of all local objectives is evaluated, and if this score is above the score of the non-mutated sequence, the candidate sequence becomes the new reference sequence. The loop stops when a variant is found for which the total objective score equals the theoretical maximum (in the case where all objectives specify a theoretical maximum), when no improvement of the sequence was observed after a number of successive iterations (this *stagnation threshold* can be set by the user, and defaults to 150), or when all variants have been evaluated, at which case the sequence variant which scored highest is retained.

##### Custom search methods

Some specifications cannot be easily met with an exhaustive or random search. For instance, assume that specification `EnforcePatternOccurence(“AarI_site”)` is used to create a AarI site (CACCTGC) in a 50bp sequence (possibly also constrained by other specifications). The size of the mutation space, 4^50^, prohibits an exhaustive search, and a random search is unlikely to create a CACCTGC site in the sequence.

Therefore, the `EnforcePatternOccurence` class of specification implements its own constraint resolution strategy, which it substitutes to the solver’s. First, an AarI site is placed in the middle of the sequence, then all constraints are evaluated and any new breach created in the operation is resolved (using an exhaustive or random search). If this resolution succeeds, then the AarI site has been successfully created in the sequence. Otherwise, the algorithm makes a new attempt, shifting the insertion location by a few nucleotides, and iterates until the site creation of the pattern in compliance with other constraints succeeds, or until all possible insertion locations have been considered, at which case it is proven that the pattern cannot be inserted in the sequence, and the constraint resolution fails with an error.

## Notes

https://github.com/Edinburgh-Genome-Foundry/DnaChisel

## References

Casini, A. et al. (2014). R2oDNA designer: Computational design of biologically neutral synthetic DNA sequences. ACS Synthetic Biology, 3(8), 525–528.

Claassens, N. J. et al. (2017). Improving heterologous membrane protein production in Escherichia coli by combining transcriptional tuning and codon usage algorithms. PLoS ONE.

Cock, P. J. A. et al. (2009). Biopython: Freely available Python tools for computational molecular biology and bioinformatics. Bioinformatics, 25(11), 1422–1423.

Guimaraes, J. C. et al. (2014). D-Tailor: Automated analysis and design of DNA sequences. Bioinformatics, 30(8), 1087–1094.

Kosuri, S. and Church, G. M. (2014). Large-scale de novo DNA synthesis: Technologies and applications. Nature Methods, 11(5), 499–507.

Nakamura, Y. (2000). Codon usage tabulated from international DNA sequence databases: status for the year 2000. Nucleic Acids Research.

Oberortner, E. et al. (2017). Streamlining the Design-to-Build Transition with Build-Optimization Software Tools. ACS Synthetic Biology, 6(3), 485–496.

Raab, D. et al. (2010). The GeneOptimizer Algorithm: Using a sliding window approach to cope with the vast sequence space in multiparameter DNA sequence optimization. Systems and Synthetic Biology, 4(3), 215–225.

Richardson, S. M. et al. (2012). Design-a-gene with genedesign. Methods in Molecular Biology, 852, 235–247.

## Bibliography

Angov, E., Hillier, C. J., Kincaid, R. L., & Lyon, J. A. (2008). Heterologous protein expression is enhanced by harmonizing the codon usage frequencies of the target gene with those of the expression host. PLoS ONE. https://doi.org/10.1371/journal.pone.0002189

Casini, A., Christodoulou, G., Freemont, P. S., Baldwin, G. S., Ellis, T. & Macdonald, J. T. (2014). R2oDNA designer: Computational design of biologically neutral synthetic DNA sequences. ACS Synthetic Biology. https://doi.org/10.1021/sb4001323

Chung BK, Lee DY. Computational codon optimization of synthetic gene for protein expression. BMC Syst Biol. 2012 https://doi.org/10.1186/1752-0509-6-134

Hale and Thompson, Codon Optimization of the Gene Encoding a Domain from Human Type 1 Neurofibromin Protein … Protein Expression and Purification 1998. https://doi.org/10.1006/prep.1997.0825

Jayaraj et. al. GeMS: an advanced software package for designing synthetic genes, Nucleic Acids Research, 2005 https://doi.org/10.1093/nar/gki614

Langmead, B., Trapnell, C., Pop, M., & Salzberg, S. L. (2009). Ultrafast and memory-efficient alignment of short DNA sequences to the human genome. Genome Biology. https://doi.org/10.1186/gb-2009-10-3-r25

Mignon, C., Mariano, N., Stadthagen, G., Lugari, A., Lagoutte, P., Donnat, S., … Werle, B. (2018). Codon harmonization – going beyond the speed limit for protein expression. FEBS Letters. https://doi.org/10.1002/1873-3468.13046

Scher, E., Cohen, S. B., & Sanguinetti, G. (2019). PartCrafter: find, generate and analyze BioParts. Synthetic Biology. https://doi.org/10.1093/synbio/ysz014

Sharp, Paul M.; Li, Wen-Hsiung (1987). The codon adaptation index-a measure of directional synonymous codon usage bias, and its potential applications. Nucleic Acids Research. 15 (3): 1281–1295. PMC 340524. PMID 3547335. https://doi.org/10.1093/nar/15.3.1281

Stormo, G. D., Schneider, T. D., Gold, L., & Ehrenfeucht, A. (1982). Use of the “perceptron” algorithm to distinguish translational initiation sites in E. coli. Nucleic Acids Research. https://doi.org/10.1093/nar/10.9.2997

Untergasser, A., Cutcutache, I., Koressaar, T., Ye, J., Faircloth, B. C., Remm, M., & Rozen, S. G. (2012). Primer3-new capabilities and interfaces. Nucleic Acids Research. https://doi.org/10.1093/nar/gks596

